# Investigating expression of a human optimized *cas9* transgene in *Neurospora crassa*

**DOI:** 10.1101/2020.12.29.424703

**Authors:** Natalie Burrell, Nicholas A. Rhoades, Amy Boyd, Jim Mierendorf, Aykhan Yusifov, Austin Harvey, Kevin Edwards, Laura Vogel, Thomas M. Hammond

**Affiliations:** School of Biological Sciences, Illinois State University, Normal, Illinois 61790

**Author notes:** These authors contributed equally to this work (Natalie Burrell, Nicholas A. Rhoades). Corresponding author: Thomas M. Hammond, School of Biological Sciences, Illinois State University, Normal, IL 61790.

**Keywords:** heterologous expression, codon optimization, translation, Cas9, fungi

## Abstract

The CRISPR-associated Cas9 enzyme is used in molecular biology to engineer the genomes of a wide range of organisms. While Cas9 can be injected or transfected into a target cell to achieve the desired goal, there are situations where stable expression of Cas9 within a target organism is preferable. Here, we show that the model filamentous fungus *Neurospora crassa* is recalcitrant to heterologous expression of a human-optimized version of *Streptococcus pyogenes cas9*. Furthermore, partial optimization of *cas9* by synonymous codon exchange failed to improve its expression in the fungus. Finally, we show that transgene expression can be detected when *cas9^Hs^* sequences are placed in the 3’ UTR regions of transgene-derived mRNAs, but not when the same sequences are in the translated part of the transgene-derived mRNA. This finding suggests that the primary obstacle to high *cas9^Hs^* expression levels in *N. crassa* is translational in nature.

## INTRODUCTION

The availability of CRISPR-associated (Cas) systems for use in genome engineering has accelerated research on organisms that were historically difficult to engineer with traditional techniques (Knott and Doudna 2018; Wang *et al*. 2020; Zhu *et al*. 2020). Cas technology has also accelerated the field of gene driver research, where it offers the possibility of engineering synthetic gene drivers for the control of disease spreading organisms (Esvelt *et al*. 2014). For these reasons, we sought to establish a robust *Streptococcus pyogenes* Cas9-based system for use in the model filamentous fungus *Neurospora crassa*.

## MATERIALS AND METHODS

### Strains and media

Strains are listed in Table 1. Vogel’s minimal medium (VMM) (Vogel 1956) with and without 2% agar was used for vegetative propagation of all strains. L-histidine was added to VMM at 500 mg / L when needed to support growth of *his-3* strains. Hygromycin B (GoldBio, H-270) was included in media at concentrations of 200 μg / ml to select for hygromycin-resistant transformants. Crosses were performed on synthetic crossing medium (pH 6.5) with 1.5 % sucrose (Westergaard and Mitchell 1947).

**Table 1.** Strains used in this study

| Strain name (alias) | Strain genotype<sup>a</sup> |
| --- | --- |
| F2-23 (RTH1005.1) | *rid; fl A* |
| F2-26 (RTH1005.2) | *rid; fl a* |
| ISU-3121 (RTH61.3.1.2) | *rid his-3; gfp-sad-6-hph a* |
| ISU-3866 (RAB1.8) | *rid A* |
| ISU-3994 (RAB22.4) | *rid his-3<sup>+</sup>::gfp-cas9<sup>Hs</sup>-hph; mus-51<sup>Δ</sup>::bar A* |
| ISU-4092 (TAB38.1) | *rid his-3<sup>+</sup>::gfp-cas9<sup>HsC991</sup>; mus-51<sup>Δ</sup>::bar a* (ISU-4145 het) |
| ISU-4107 (TAB44.1) | *rid his-3<sup>+</sup>::gfp-cas9<sup>HsC318</sup>; mus-51<sup>Δ</sup>::bar a* (ISU-4145 het) |
| ISU-4145 (HAB11.53.2) | *rid his-3<sup>+</sup>::tef1(p)-cas9<sup>Hs</sup>; mus-51<sup>Δ</sup>::bar a* |
| ISU-4231 (RAB20.1) | *rid his-3<sup>+</sup>::gfp-cas9<sup>HsC4</sup>-hph a* |
| ISU-4232 (HAB45.1.1) | *rid his-3<sup>+</sup>::gfp-cas9<sup>HsC213</sup>; mus-51<sup>Δ</sup>::bar a* |
| ISU-4233 (RAB1.7) | *rid a* |
| ISU-4234 (HPM32.14.2) | *rid his-3<sup>+</sup>::gfp-cas9<sup>HsC135</sup>; mus-51<sup>Δ</sup>::bar a* |
| ISU-4235 (HPM33.11.2) | *rid his-3<sup>+</sup>::gfp-cas9<sup>HsC70</sup> a* |
| ISU-4236 (HPM34.1.1) | *rid his-3<sup>+</sup>::gfp-cas9<sup>HsC25</sup> a* |
| ISU-4522 (RAY25.1.4) | *rid his-3<sup>+</sup>::gfp-cas9<sup>HsC62</sup> a* |
| ISU-4618 (RAY11.13) | *rid his-3<sup>+</sup>::gfp-cas9<sup>HsC60</sup>; mus-51<sup>Δ</sup>::bar a* |
| ISU-4624 (RAY13.1) | *rid his-3<sup>+</sup>::gfp-cas9<sup>HsC40</sup> a* |
| ISU-4875 (CTH293.3.3) | *rid his-3<sup>+</sup>::gfp-stop-cas9<sup>HsC213</sup>; mus-51<sup>Δ</sup>::bar a* |
| ISU-4884 (CNR307.3.1) | *rid his-3<sup>+</sup>::gfp-cas9<sup>NcC213</sup>; mus-52<sup>Δ</sup>::bar A* |
| ISU-4886 (CNR308.4.1) | *rid his-3<sup>+</sup>::gfp-cas9<sup>NcC60</sup>; mus-52<sup>Δ</sup>::bar A* |
| ISU-4888 (RABOB.36.4.1) | *rid his-3<sup>+</sup>::gfp-cas9<sup>HsC1209</sup>; mus-51<sup>Δ</sup>::bar a* |
| ISU-4889 (RABLG.37.1.2) | *rid his-3<sup>+</sup>::gfp-cas9<sup>HsC1078</sup>; mus-51<sup>Δ</sup>::bar a* |
| ISU-4892 (RABBL.39.1.1) | *rid his-3<sup>+</sup>::gfp-cas9<sup>HsC888</sup>; mus-51<sup>Δ</sup>::bar a* |
| ISU-4894 (RABJE.41.1.1) | *rid his-3<sup>+</sup>::gfp-cas9<sup>HsC656</sup>; mus-51<sup>Δ</sup>::bar a* |
| ISU-4895 (RABTrH.42.1.1) | *rid his-3<sup>+</sup>::gfp-cas9<sup>HsC535</sup>; mat a* |
| ISU-4897 (RAB.XL.43.2.2) | *rid his-3<sup>+</sup>::gfp-cas9<sup>HsC443</sup>; mat a* |
| ISU-4933 (CNR326.2.1/76.2.1) | *rid his-3<sup>+</sup>::gfp-stop-cas9<sup>Hs</sup>; mus-51<sup>Δ</sup>::bar a* |
| ISU-4934 (CNR327.1.1/77.1.1) | *rid his-3<sup>+</sup>::gfp-stop-cas9<sup>HsC4</sup>; mus-51<sup>Δ</sup>::bar a* |
| ISU-4935 (CNR336.1.1/104.1.1) | *rid his-3<sup>+</sup>::gfp-stop-cas9<sup>HsC213</sup>; mus-51<sup>Δ</sup>::bar a* |
| ISU-4936 (CNR337.1.1/124.1.1) | *rid his-3<sup>+</sup>::gfp-stop-cas9<sup>HsC135</sup>; mus-51<sup>Δ</sup>::bar a* |
| ISU-4937 (CNR339.1.1/126.1.1) | *rid his-3<sup>+</sup>::gfp-stop-cas9<sup>HsC25</sup>; mus-51<sup>Δ</sup>::bar a* |
| ISU-4938 (CNR347.1.5/v193h.5) | *rid his-3<sup>+</sup>::gfp-stop-cas9<sup>HsC62</sup>; mus-51<sup>Δ</sup>::bar a* |
| ISU-4939 (CNR340.2.1/161.2.1) | *rid his-3<sup>+</sup>::gfp-stop-cas9<sup>HsC60</sup>; mus-51<sup>Δ</sup>::bar a* |
| ISU-4941 (CNR341.1.2 /163.1.2) | *rid his-3<sup>+</sup>::gfp-stop-cas9<sup>HsC40</sup>; mus-51<sup>Δ</sup>::bar a* |
| ISU-4942 (CNR328.1.1/95.1.1) | *rid his-3<sup>+</sup>::gfp-stop-cas9<sup>HsC1209</sup>; mus-51<sup>Δ</sup>::bar a* |
| ISU-4943 (CNR329.1.1/96.1.1) | *rid his-3<sup>+</sup>::gfp-stop-cas9<sup>HsC1078</sup>; mus-51<sup>Δ</sup>::bar a* |
| ISU-4944 (CNR330.1.1/97.1.1) | *rid his-3<sup>+</sup>::gfp-stop-cas9<sup>HsC991</sup>; mus-51<sup>Δ</sup>::bar a* |
| ISU-4945 (CNR331.1.1/98.1.1) | *rid his-3<sup>+</sup>::gfp-stop-cas9<sup>HsC888</sup>; mus-51<sup>Δ</sup>::bar a* |
| ISU-4946 (CNR332.1.1/v100.1.1) | *rid his-3<sup>+</sup>::gfp-stop-cas9<sup>HsC656</sup>; mus-51<sup>Δ</sup>::bar a* |
| ISU-4947 (CNR333.2.1/101.2.1) | *rid his-3<sup>+</sup>::gfp-stop-cas9<sup>HsC535</sup>; mus-51<sup>Δ</sup>::bar a* |
| ISU-4948 (CNR334.1.2/102.1.2) | *rid his-3<sup>+</sup>::gfp-stop-cas9<sup>HsC443</sup>; mus-51<sup>Δ</sup>::bar a* |
| ISU-4949 (CNR335.2.1/103.2.1) | *rid his-3<sup>+</sup>::gfp-stop-cas9<sup>HsC318</sup>; mus-51<sup>Δ</sup>::bar a* |
| ISU-4951 (CNR348.1.7/307.1.7) | *rid his-3<sup>+</sup>::gfp-stop-cas9<sup>HsC796</sup>; mus-51<sup>Δ</sup>::bar a* |
| ISU-4952 (CNR349.1.2/308.1.2) | *rid his-3<sup>+</sup>::gfp-stop-cas9<sup>HsC714</sup>; mus-51<sup>Δ</sup>::bar a* |
| ISU-4953 (CNR350.2.8/309.2.8) | *rid his-3<sup>+</sup>::gfp-stop-cas9<sup>HsC612</sup>; mus-51<sup>Δ</sup>::bar a* |
| P8-42 | *rid his-3; mus-51<sup>Δ</sup>::bar a* |
| P8-43 | *rid his-3; mus-52<sup>Δ</sup>::bar A* |

<sup>a</sup>The symbols *cas9<sup>Hs</sup>* and *cas9<sup>Nc</sup>* are used for human and *N. crassa* optimized Cas9 coding sequences, respectively.

### *N. crassa* transformation

Transformation of *N. crassa* was performed as described by Rhoades et al. (2019b), except that the recovery step was omitted when transformants were selected for histidine prototrophy. *N. crassa* strain P8-42 was transformed with pAH41.4, a plasmid containing *tef-1(p)-cas9^Hs^*. Transformants were selected by their ability to grow on medium lacking histidine.

Transformation vectors v76, v77, v95, v96, v97, v98, v100, v101, v102, v103, v104, v124, v125, v126, v161, v163, and v193, which convert a *tef-1(P)-cas9^Hs^* transgene to a *ccg-1(P)-gfp-cas9^Hs^* transgene, were transformed into strain ISU-4145. Transformants were selected for resistance to hygromycin B. Transformation vectors v76-stop, v77-stop, v95-stop, v96-stop, v97-stop, v98-stop, v100-stop, v101-stop, v102-stop, v103-stop, v104-stop, v124-stop, v126-stop, v161-stop, v163-stop, v193-stop, v296-stop, v307-stop, v308-stop, and v309-stop, which convert a *tef-1(P)-cas9^Hs^* transgene to a *ccg-1(P)-gfp-stop-cas9^Hs^* transgene, were transformed into strain ISU-4145. Transformants were selected for resistance to hygromycin B. Plasmids pNR206.2 and pNR207.1 contain *gfp-cas9^NcC213^* and *gfp-cas9^NcC60^* transgenes, respectively, and these plasmids were used to transform strain P8-43. Transformants were selected for histidine prototrophy. Homokaryotic strains were isolated from transformants by single spore purification (from conidium or ascospores) and confirmed to be homokaryotic with polymerase chain reaction (PCR)-based genotyping assays.

### Transgene construction and insertion

*tef-1(P)-cas9^Hs^*: A *tef-1(P)-cas9^Hs^*-containing plasmid (pAH41.4) was constructed by amplifying the *N. crassa tef-1* promoter region (*tef-1[P]*) from *N. crassa* genomic DNA with primers 598 and 599 by PCR and inserting the PCR product into the *Spe*I site of Plasmid 43802 (Addgene, DiCarlo *et al*. 2013). The resulting plasmid was used as the template for PCR-based amplification of *tef-1(P)-cas9^Hs^* with primers 610 and 589. The PCR product was inserted into the *Not*I site of the *his-3*-targeting plasmid pTH1150.11 (GenBank MN872812.1) to produce plasmid pAH41.4. Primer sequences are listed in Table S1. Sequences of *N. crassa* genes, such as *tef-1* and the *tef-1* promoter region, were obtained from FungiDB (Stajich *et al*. 2012). *gfp-cas9^Hs^*: The *gfp-cas9^Hs^* and *gfp-stop-cas9^Hs^* transgenes, including all truncated versions (e.g., *gfp-cas9^HsC1209^, gfp-slop-cas9^HsC1078^*. etc.) were constructed by double-joint PCR (DJ-PCR) (Hammond *et al*. 2011) with primers and templates described in Tables S1 and S2.

*gfp-cas9^NcC213^*: Primers 2021 and 2022 were used to amplify a single PCR product containing *hph, ccg-1(P)* (McNally and Free 1988), *gfp*, and a *(GA)_5_*-linker from plasmid pTH1117.12 (GenBank JF749202.1). Primers 1980 and 624 were used to amplify a single PCR product containing *cas9^Nc213^* from pNR177.2 (described below). The two products were fused by PCR, and the fusion product was used as a template for amplification with primers 2024 and 2025. The amplified product was digested with *Not*I and cloned into the *Not*I site of the *his-3*-targeting plasmid pTH1150.11 to create plasmid pNR206.2.

*gfp-cas9^NcC60^*: The *gfp-cas9^NcC60^* transgene was assembled similarly to the *gfp-cas9^NcC213^* transgene except that primer 1980 was exchanged for primer 1983. The final amplification product was cloned into the *Not*I site of pTH1150.11 to create plasmid pNR207.1.

### Codon optimization

We used a publicly available RNAseq dataset (SRR1055991, Wu *et al*. 2014) to identify 100 of the most highly expressed protein-coding genes in the *N. crassa* nuclear genome (Table S3) by aligning RNA sequences to all predicted *N. crassa* protein coding genes as described in Samarajeewa et al. (2017). Genes with the highest “reads per kilobase exon model” (RPKM) values (Mortazavi *et al*. 2008) were considered to be the most highly expressed. Relative adaptiveness (RA) values (Sharp and Li 1987) were then calculated for each codon (Table S4). RA values were calculated by dividing the observed frequency of each codon in the set of 100 highly expressed genes by the frequency of the most common synonymous codon in the same set of genes. Codon optimization of *cas9* for expression in *N. crassa* was then performed by replacing *cas9^Hs^* codons in *cas9^Hs^* as follows: 1) the synonymous codon with the highest RA value was used for all occurrences of amino acids C, E, F, H, I, K, L, M, N, Q, W, and Y, as well as the stop codon; 2) the synonymous codon with the second highest RA value was used for all occurrences of A, D, G, P, and R; and, 3) a combination of synonymous codons with either the first or second highest RA values was used for amino acids S, T, and V. Use of the second most optimal codon for some amino acids helped reduce the overall GC content of the optimized sequence. Reducing GC content allowed the optimized sequence to be synthesized as a gBlock^®^ (Integrated DNA technologies). A gBlock^®^ DNA fragment containing the optimized *cas9^Nc^* sequence (Figure S1) was cloned to pJET1.2 to create pNR177.2.

### Visualization of GFP in conidia by fluorescence microscopy

Fluorescence microscopy and imaging was performed with a Leica DMBRE microscope or a Leica SP8 confocal microscope. For imaging with a Leica DMBRE system, *N. crassa* cultures were incubated on VMM for 1-2 days at 32°C followed by 2-4 days at room temperature. Conidial suspensions were placed on a standard microscope slide for imaging. A 40× objective was used. GFP-signal was collected with a 20 second exposure for all strains. Cropped raw images were used without additional modifications. For imaging with the Leica SP8 confocal microscope, the same growth conditions were followed as stated for imaging with the Leica DMBRE system. Conidial suspensions were prepared, and then incubated in a 37°C shaker for 2 hours before imaging. Conidial suspensions were placed on a standard microscope slide for imaging. All the images were acquired with the same settings so that they could be directly compared. A 63 ×/1.40 oil objective was used, with a white light laser set to a wavelength of 488 nm. Fluorescence was detected between the wavelengths of 500–560nm, with 8× line averaging. Images were assembled in Photoshop; GFP is shown with original contrast.

### Quantification of GFP in conidia by flow cytometry

Fresh *N. crassa* cultures were prepared by qualitative transfer of conidia to 2.5 ml of solid VMM slants in 16×100 mm glass culture tubes. The inoculated culture tubes were incubated for two days in a 32°C incubator and 5–6 days at room temperature on a laboratory bench top. Sterile wood applicators were used to transfer conidia to 1× PBS (137 mM NaCl, 2,7 mM KCl, 10mM Na_2_HPO_4_, 1.8 mM KH_2_PO_4_, pH 7.4) (Figure S2). Conidial suspensions were then analyzed with a BD FACSMelody system (equipped with 488 nm, 561 nm, and 640 nm lasers) and FACSChorus software. Conidia were gated on FSC and SSC and MFI of GFP histograms were analyzed using a 1.5 neutral density filter. Raw MFI values are provided in Table S5.

## RESULTS

### A *cas9^Hs^* transgene is poorly expressed in *N. crassa*

We began our studies by constructing a *cas9^Hs^* transgene (Fig. 1A) and inserting it downstream of *his-3* on chromosome I (Fig. 1B). We then constructed a second transgene, *gfp-cas9^Hs^*, to express a GFP-Cas9 fusion protein (Fig. 1, C–E). This second transgene was constructed to allow us to determine if Cas9, which includes an SV40 nuclear localization signal on its C-terminus, localizes to the *N. crassa* nucleus. However, when we examined the *gfp-cas9^Hs^* strain by fluorescence microscopy, we failed to detect a GFP signal (Fig 2, A–B). Because we focused our assays on *N. crassa* conidia (asexual spores), we considered the possibility that the promoter used to drive expression of *gfp-cas9^Hs^ (ccg-1[P])* was insufficient for expression of the fusion protein in this cell type. Thus, as a control, we examined conidia from a *gfp-sad-6* transgene-carrying strain. SAD-6 is a meiotic silencing by unpaired DNA (MSUD) protein (Samarajeewa *et al*. 2014) and it was chosen as a control for Cas9 expression because it is larger than Cas9 (210 kD vs 159 kD) and because, like the *gfp-cas9^Hs^* transgene, the *gfp-sad-6* transgene is driven by the *ccg-1* promoter. In contrast to GFP-Cas9, GFP-SAD-6 was easily detected by fluorescence microscopy (Figure S3, compare A and B). These observations suggest that the *ccg-1* promoter should be sufficient for expression of GFP-Cas9 in *N. crassa* conidia.

**Figure 1.**
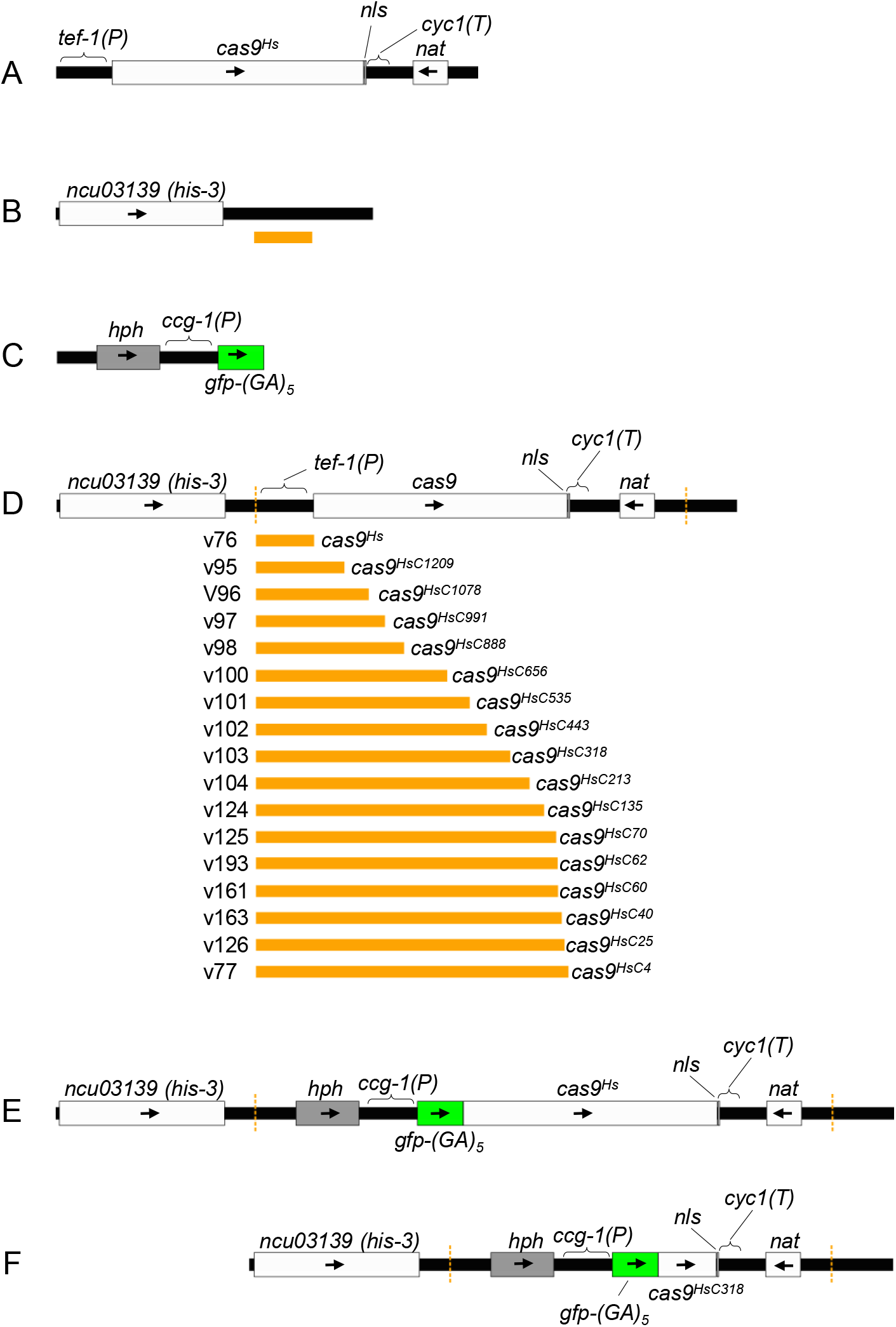
Transgene illustrations. (A) Diagram of the *tef-1(P)::cas9^Hs^* transgene. The *cas9^Hs^* transgene contains the *N. crassa tef-1* promoter (*tef-1[P]*), the *cas9^Hs^*-coding sequence, an SV40 nuclear localization signal (*nls*), an *S. cerevisiae cyc1* terminator (*cyc1[T]*), and a nourseothricin resistance cassette (*nat*). (B) Diagram of the *N. crassa his-3* locus. The orange horizontal bar marks the DNA interval that was deleted and replaced with the *tef-1(P)::cas9^Hs^* transgene. The diagram is drawn to scale, and the orange bar represents a length of 919 bp. The *his-3* coding region is depicted by the white rectangle. (C) Diagram of the *ccg-1(P)::gfp-(GA)5* construct used to construct *gfp-cas9^Hs^* transgenes. Note that the *gfp* sequence in every *gfp*-containing transgene examined in this study is immediately followed by the coding sequence for a ten amino acid glycine-alanine linker ([GA]_5_). (D) Diagram showing the DNA intervals of *tef-1(P)::cas9^Hs^* that were deleted and replaced with *ccg-1(P)::gfp*. The orange vertical dashed lines mark the two *tef-1(P)::cas9^Hs^* transgene borders. The orange horizontal bars mark the intervals that were deleted and replaced by *ccg-1(P)::gfp* with various transformation vectors (e.g., v76, v95, etc.). (E) Diagram of the *ccg-1(P)::gfp-cas9^Hs^* transgene obtained by transformation of strain ISU-4145 with transformation vector v76. (F) Diagram of the *ccg-1(P)::gfp-cas9^HsC318^* transgene obtained by transformation of strain ISU-4145 with transformation vector v103.

**Figure 2.**
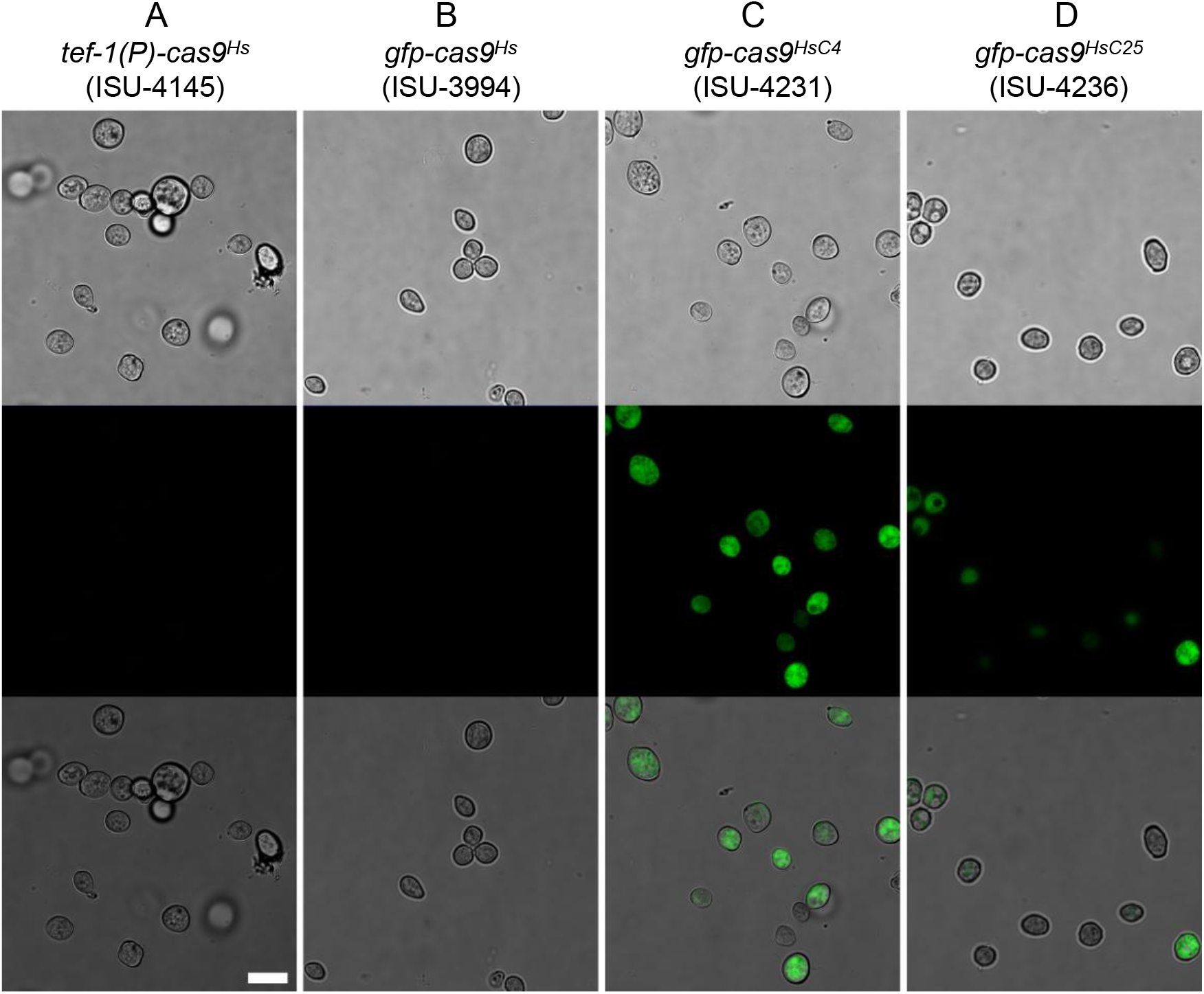
Confocal microcopy-based analysis of transgene expression. (A–D) Conidia from a *tef-1(P)-cas9^Hs^* strain (ISU-4145), a *gfp-cas9^Hs^* strain (ISU-3994), a *gfp-cas9^HsC4^* strain (ISU-4231), and a *gfp-cas9^HsC25^* strain (ISU-4236) were examined by confocal microscopy. Upper panels, transmitted light image; middle panels, GFP signal; lower panels, overlay of transmitted and GFP images. The white horizontal bar represents 10 μm.

### Partial codon optimization of *cas9* for expression in *N. crassa*

The above results suggest that *cas9^Hs^* sequences are poorly expressed in *N. crassa*. To identify a segment of *cas9^Hs^* that could be expressed more robustly in the organism, and concomitantly identify regions of *cas9^Hs^* that prevent its robust expression, we dissected the *cas9^Hs^* coding region with a series of *gfp-cas9^Hs<u>C#</u>^* transgenes (Fig. 1D). “C#” specifies the number of Cas9 amino acids in the GFP-tagged and N-terminally truncated Cas9 protein (for example, *cas9^HsC60^* encodes only the C-terminal 60 amino acids of Cas9. The number “60” does not include the seven amino acids of the C-terminal NLS). We constructed 16 of these *gfp-cas9^HsC#^* transgenes and examined GFP levels in conidia from the *gfp-cas9^HsC#^* transgene-carrying strains by flow cytometry. Interestingly, we found that all transgenes containing more than 212 *cas9^Hs^* codons failed to produce a GFP signal at detectable levels and that GFP levels increased rapidly as the number of *cas9^Hs^* codons approached zero (Fig 2, C and D; Fig. 3A).

**Figure 3.**
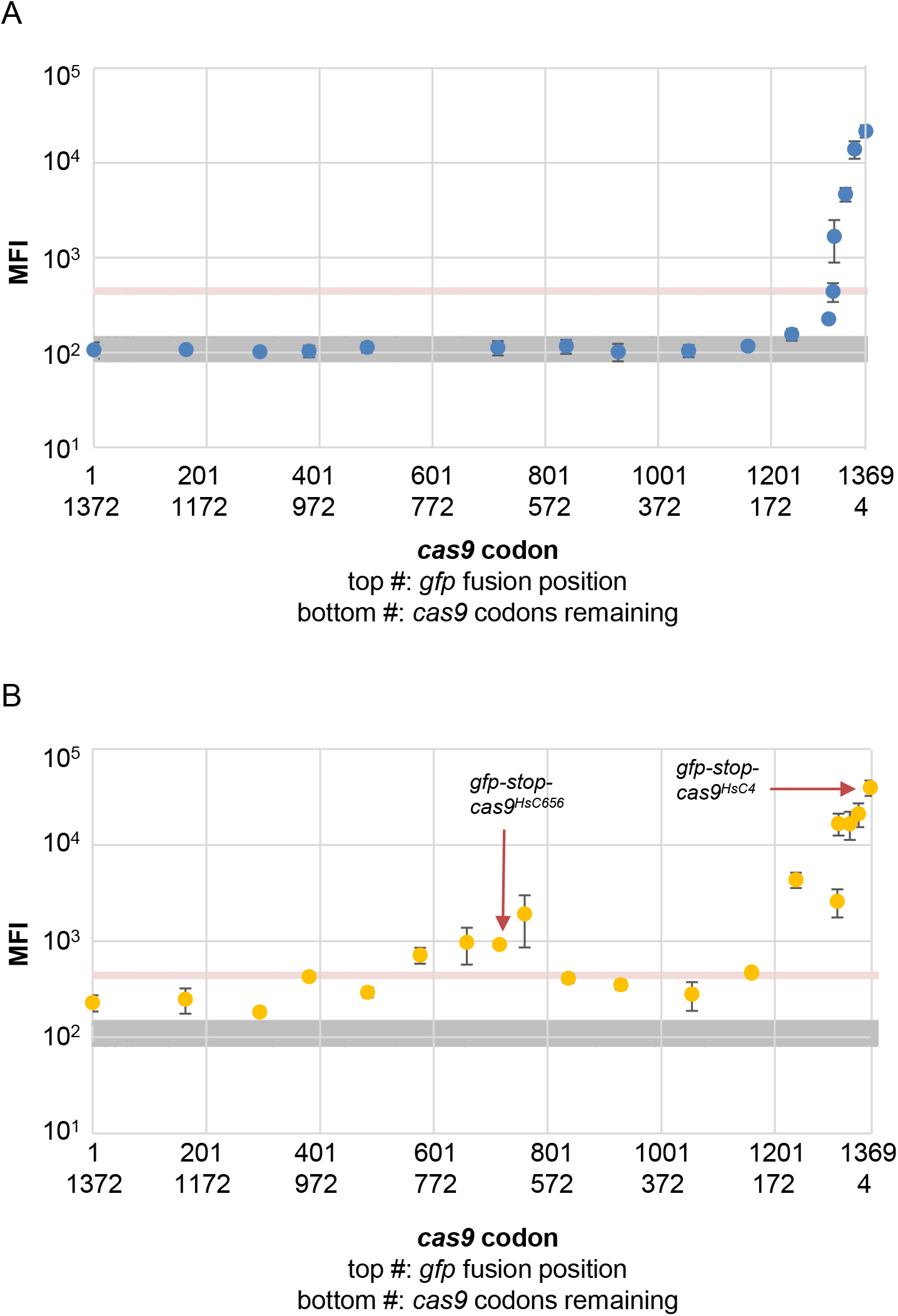
Flow cytometry-based analysis of transgene expression. (A) Conidia from *gfp-cas9^Hs^* and *gfp-cas9^HsC#^* strains were analyzed for the presence of GFP by flow cytometry. Mean fluorescence intensity (MFI) values are plotted on the Y axis relative to the *cas9^Hs^* codon number to which *gfp* is fused in each transgene. Note that *cas9^Hs^* contains 1380 codons, including seven codons for the SV40 NLS and one codon for the translational stop signal). The gray shaded region of the chart marks the range of MFI values (+/- standard deviation) obtained for negative control strains. The pink shaded region marks the MFI value (+/- standard deviation) obtained for the *gfp-sad-6* positive control strain. MFI values are averages obtained from two assays that were completed one day apart. Error bars are standard deviation values. The data used to construct the chart are provided in Table 2. Raw data is provided in Table S5. (B) Conidia from *gfp-stop-cas9^Hs^* and *gfp-stop-cas9^HsC#^* strains were analyzed for the presence of GFP by flow cytometry as in panel A. MFI values are averages of values obtained from two assays that were completed one day apart. The data used to construct the chart is provided in Table 3. Raw data is provided in Table S5.

**Table 2.** *gfp-cas9^Hs^* expression levels

| Strain name | Transgene name | vector | *gfp*<br>seq<sup>a</sup> | *cas9*<br>seq<sup>b</sup> | Average<br>MFI<sup>c</sup> | stdev |
| --- | --- | --- | --- | --- | --- | --- |
| **Control strains** |  |  |  |  |  |  |
| ISU-4233 | *none* | na | na | na | 107.30 | 6.78 |
| ISU-3866 | *none* | na | na | na | 101.15 | 5.32 |
| ISU-4145 | *cas9<sup>Hs</sup>* | na | na | no<sup>d</sup> | 113.12 | 28.33 |
| ISU-3121 | *gfp-sad-6* | na | yes | na | 445.07 | 13.30 |
| **Test strains** |  |  |  |  |  |  |
| ISU-3994 | *gfp-cas9<sup>Hs</sup>* | v76 | no | no | 107.22 | 21.42 |
| ISU-4888 | *gfp-cas9<sup>Hs</sup>C1209* | v95 | yes | no | 107.82 | 0.56 |
| ISU-4889 | *gfp-cas9<sup>Hs</sup>C1078* | v96 | yes | no | 102.49 | 7.25 |
| ISU-4092 | *gfp-cas9<sup>Hs</sup>C991* | v97 | no | no | 103.45 | 14.27 |
| ISU-4892 | *gfp-cas9<sup>Hs</sup>C888* | v98 | yes | no | 113.74 | 13.53 |
| ISU-4894 | *gfp-cas9<sup>Hs</sup>C656* | v100 | yes | no | 112.94 | 19.64 |
| ISU-4895 | *gfp-cas9<sup>Hs</sup>C535* | v101 | yes | no | 116.87 | 19.95 |
| ISU-4897 | *gfp-cas9<sup>Hs</sup>C443* | v102 | yes | no | 101.96 | 21.30 |
| ISU-4107 | *gfp-cas9<sup>Hs</sup>C318* | v103 | no | no | 104.77 | 15.48 |
| ISU-4232 | *gfp-cas9<sup>Hs</sup>C213* | v104 | yes | yes | 116.96 | 11.79 |
| ISU-4234 | *gfp-cas9<sup>Hs</sup>C135* | v124 | yes | yes | 155.74 | 21.98 |
| ISU-4235 | *gfp-cas9<sup>Hs</sup>C70* | v125 | yes | yes | 227.29 | 10.30 |
| ISU-4522 | *gfp-cas9<sup>Hs</sup>C62* | v193 | yes | yes | 440.88 | 99.47 |
| ISU-4618 | *gfp-cas9<sup>Hs</sup>C60* | v161 | yes | yes | 1686.60 | 799.05 |
| ISU-4624 | *gfp-cas9<sup>Hs</sup>C40* | v163 | yes | yes | 4674.03 | 783.60 |
| ISU-4236 | *gfp-cas9<sup>Hs</sup>C25* | v126 | yes | yes | 13973.04 | 2898.18 |
| ISU-4231 | *gfp-cas9<sup>Hs</sup>C4* | v77 | yes | yes | 21681.28 | 3130.25 |
| ISU-4884 | *gfp-cas9<sup>Nc</sup>C213* | v293 | yes | yes | 97.65 | 4.33 |
| ISU-4886 | *gfp-cas9<sup>Nc</sup>C60* | v294 | yes | yes | 519.11 | 116.86 |

<sup>a</sup> “yes” indicates that the *gfp* coding sequence and the ten amino acid GAGAGAGAGA linker of the *gfp-cas9* transgene were confirmed to be free of mutations in the indicated strain by Sanger sequencing. “no” means the sequence is assumed to be correct but it was not confirmed by Sanger sequencing.
<sup>b</sup> “yes” indicates that the *cas9* coding sequence was confirmed to be free of mutations in the indicated strain by Sanger sequencing. “no” means the sequence is assumed to be correct but it was not confirmed by Sanger sequencing.
<sup>c</sup> Average mean fluorescence intensity (MFI) values were calculated from MFI measurements of the same strain taken on different days (replicate assays separated in time by one day).
<sup>d</sup> The *cas9<sup>Hs</sup>* coding sequence in plasmid pAH41.1 was confirmed to be free of mutations by Sanger sequencing but the *cas9<sup>Hs</sup>* coding sequence was not confirmed to be mutation free after integration into the *N. crassa* genome.
Standard deviation values (stdev) are provided. na, not applicable.

We considered the possibility that robust expression of *cas9^Hs^* in *N. crassa* is restricted by the organism’s synonymous codon biases. To test this hypothesis, we altered the *cas9* coding sequence in the *gfp-cas9^HsC213^* and *gfp-cas9^HsC60^* transgenes to produce equivalent *gfp-cas9^NcC213^* and *gfp-cas9^NcC60^* transgenes (Figure S3). Specifically, we replaced codons that are found at low frequency in highly expressed *N. crassa* mRNAs with synonymous codons that are found at high frequency in *N. crassa* mRNAs (see methods). Interestingly, despite replacing 120 of 220 codons to make *cas9^NcC213^* (213 *cas9^Hs^* codons, 7 *nls* codons, and 1 stop codon) and replacing 40 of 68 codons to make *cas9^NcC60^* (60 *cas9^Hs^* codons, 7 *nls* codons, and 1 stop codon), neither transgene produced more GFP fusion protein than their human-optimized counterparts (Fig. 4).

**Figure 4.**
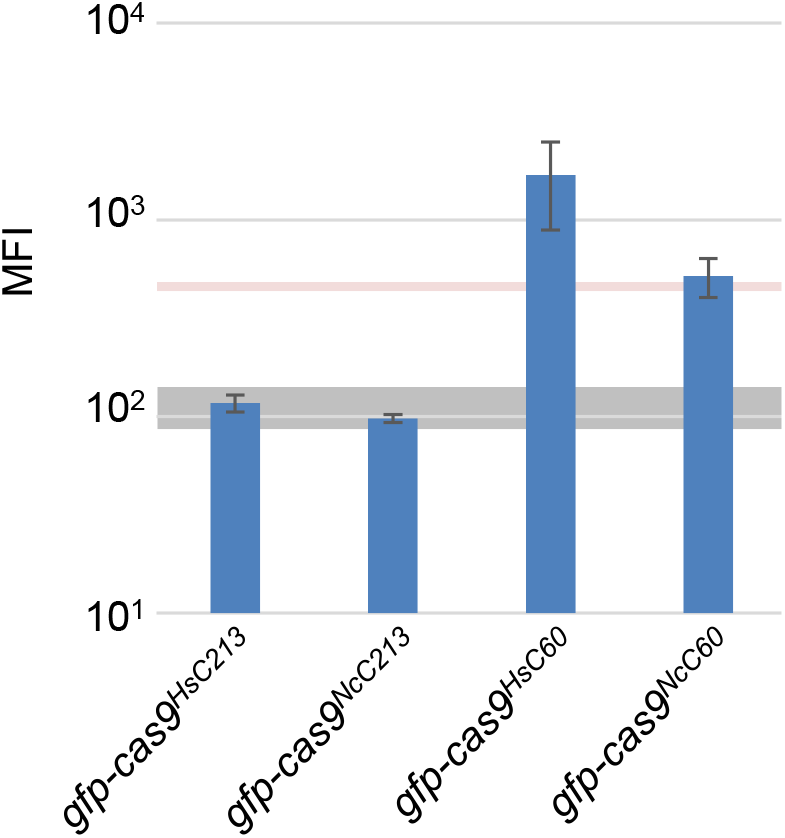
Expression profiles of similar *gfp-cas9^HsC#^* and *gfp-cas9^NcC#^* transgenes. Conidia from *gfp-cas9^HsC213^, gfp-cas9^NcC213^, gfp-cas9^HsC60^*, and *gfp-cas9^NcC60^* were analyzed for the presence of GFP by flow cytometry. The gray shaded region of the chart marks the range of MFI values (+/- standard deviation) obtained for negative control strains. The pink shaded region denotes the MFI values and standard deviation obtained for the *gfp-sad-6* positive control strain. The data used to construct the chart is provided in Table 2. Raw data is provided in Table S5.

### Translation of *cas9^Hs^* sequences negatively influences expression of *gfp-cas9^Hs^* transgenes

The tendency of GFP levels to increase as the length of *cas9^Hs^*-coding sequence decreases suggests that low GFP levels are due to problems encountered during translation of *cas9^Hs^* sequences by *N. crassa* ribosomes. To test this hypothesis, we constructed a set of *gfp-<u>stop</u>-cas9^HsC#^* transgenes. These transgenes are identical to the *gfp-cas9^HsC#^* transgenes except that they contain a stop codon between the *gfp* and *cas9^Hs^* coding sequences. Interestingly, we found that the placement of a stop codon between the *gfp* and *cas9^Hs^* coding sequences increased the level of GFP produced by all transgenes, but not in a completely uniform manner (Figure 3B and Table 3). For example, GFP levels appeared to increase around two maxima, one represented by the *gfp-stop-cas9^HsC656^* transgene and the other by the *gfp-stop-cas9^HsC4^* transgene (Figure 3B).

**Table 3.**
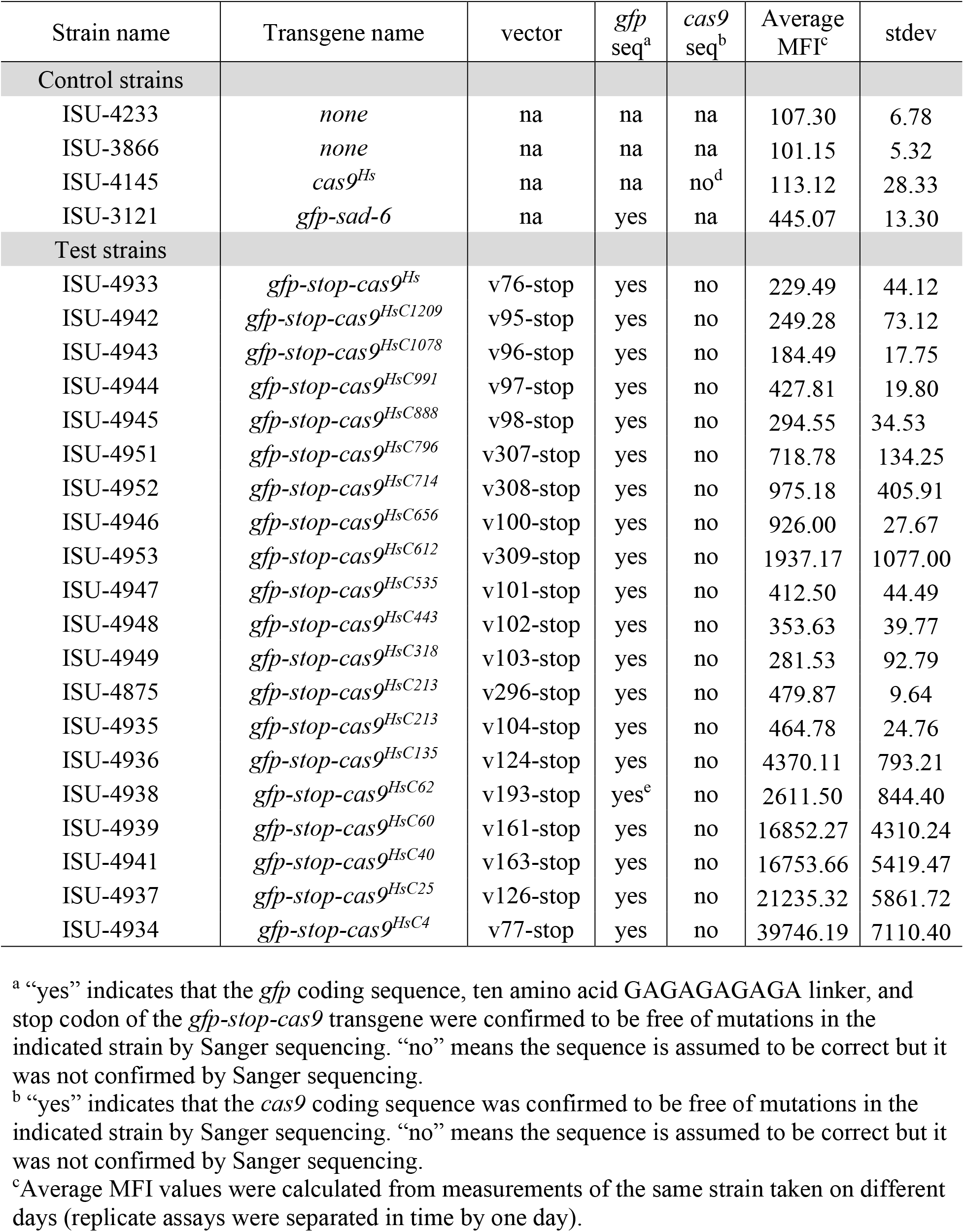
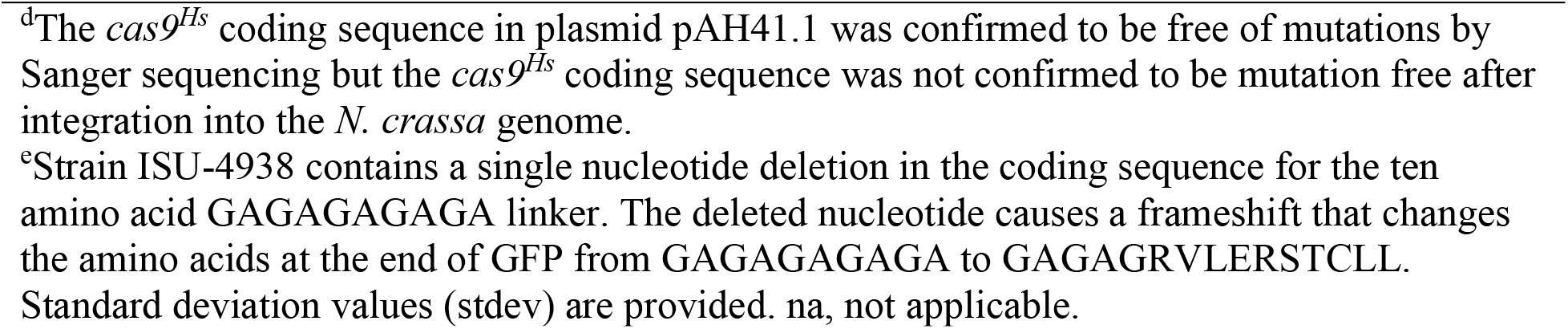
*gfp-stop-cas9^Hs^* expression levels

| Strain name | Transgene name | vector | *gfp*<br>seq<sup>a</sup> | *cas9*<br>seq<sup>b</sup> | Average<br>MFI<sup>c</sup> | stdev |
| --- | --- | --- | --- | --- | --- | --- |
| Control strains |  |  |  |  |  |  |
| ISU-4233 | *none* | na | na | na | 107.30 | 6.78 |
| ISU-3866 | *none* | na | na | na | 101.15 | 5.32 |
| ISU-4145 | *cas9<sup>Hs</sup>* | na | na | no<sup>d</sup> | 113.12 | 28.33 |
| ISU-3121 | *gfp-sad-6* | na | yes | na | 445.07 | 13.30 |
| Test strains |  |  |  |  |  |  |
| ISU-4933 | *gfp-stop-cas9<sup>Hs</sup>* | v76-stop | yes | no | 229.49 | 44.12 |
| ISU-4942 | *gfp-stop-cas9<sup>HsC1209</sup>* | v95-stop | yes | no | 249.28 | 73.12 |
| ISU-4943 | *gfp-stop-cas9<sup>HsC1078</sup>* | v96-stop | yes | no | 184.49 | 17.75 |
| ISU-4944 | *gfp-stop-cas9<sup>HsC991</sup>* | v97-stop | yes | no | 427.81 | 19.80 |
| ISU-4945 | *gfp-stop-cas9<sup>HsC888</sup>* | v98-stop | yes | no | 294.55 | 34.53 |
| ISU-4951 | *gfp-stop-cas9<sup>HsC796</sup>* | v307-stop | yes | no | 718.78 | 134.25 |
| ISU-4952 | *gfp-stop-cas9<sup>HsC714</sup>* | v308-stop | yes | no | 975.18 | 405.91 |
| ISU-4946 | *gfp-stop-cas9<sup>HsC656</sup>* | v100-stop | yes | no | 926.00 | 27.67 |
| ISU-4953 | *gfp-stop-cas9<sup>HsC612</sup>* | v309-stop | yes | no | 1937.17 | 1077.00 |
| ISU-4947 | *gfp-stop-cas9<sup>HsC535</sup>* | v101-stop | yes | no | 412.50 | 44.49 |
| ISU-4948 | *gfp-stop-cas9<sup>HsC443</sup>* | v102-stop | yes | no | 353.63 | 39.77 |
| ISU-4949 | *gfp-stop-cas9<sup>HsC318</sup>* | v103-stop | yes | no | 281.53 | 92.79 |
| ISU-4875 | *gfp-stop-cas9<sup>HsC213</sup>* | v296-stop | yes | no | 479.87 | 9.64 |
| ISU-4935 | *gfp-stop-cas9<sup>HsC213</sup>* | v104-stop | yes | no | 464.78 | 24.76 |
| ISU-4936 | *gfp-stop-cas9<sup>HsC135</sup>* | v124-stop | yes | no | 4370.11 | 793.21 |
| ISU-4938 | *gfp-stop-cas9<sup>HsC62</sup>* | v193-stop | yes<sup>e</sup> | no | 2611.50 | 844.40 |
| ISU-4939 | *gfp-stop-cas9<sup>HsC60</sup>* | v161-stop | yes | no | 16852.27 | 4310.24 |
| ISU-4941 | *gfp-stop-cas9<sup>HsC40</sup>* | v163-stop | yes | no | 16753.66 | 5419.47 |
| ISU-4937 | *gfp-stop-cas9<sup>HsC25</sup>* | v126-stop | yes | no | 21235.32 | 5861.72 |
| ISU-4934 | *gfp-stop-cas9<sup>HsC4</sup>* | v77-stop | yes | no | 39746.19 | 7110.40 |

<sup>a</sup> “yes” indicates that the *gfp* coding sequence, ten amino acid GAGAGAGAGA linker, and stop codon of the *gfp-stop-cas9* transgene were confirmed to be free of mutations in the indicated strain by Sanger sequencing. “no” means the sequence is assumed to be correct but it was not confirmed by Sanger sequencing.
<sup>b</sup> “yes” indicates that the *cas9* coding sequence was confirmed to be free of mutations in the indicated strain by Sanger sequencing. “no” means the sequence is assumed to be correct but it was not confirmed by Sanger sequencing.
<sup>c</sup> Average MFI values were calculated from measurements of the same strain taken on different days (replicate assays were separated in time by one day).
<sup>d</sup>The *cas9<sup>Hs</sup>* coding sequence in plasmid pAH41.1 was confirmed to be free of mutations by Sanger sequencing but the *cas9<sup>Hs</sup>* coding sequence was not confirmed to be mutation free after integration into the *N. crassa* genome.
<sup>e</sup>Strain ISU-4938 contains a single nucleotide deletion in the coding sequence for the ten amino acid GAGAGAGAGA linker. The deleted nucleotide causes a frameshift that changes the amino acids at the end of GFP from GAGAGAGAGA to GAGAGRVLERSTCLL. Standard deviation values (stdev) are provided. na, not applicable.

## DISCUSSION

In this study, we have presented evidence demonstrating that a human-optimized *cas9^Hs^* transgene is poorly expressed in *N. crassa*. Our major findings are as follows: 1) a *gfp-cas9^Hs^* transgene is poorly expressed relative to a control transgene *gfp-sad-6*, 2) removing the majority of *cas9^Hs^* coding sequences from the *gfp-cas9^Hs^* transgene (e.g., with *gfp-cas9^Hs**C**#^* transgenes) improves expression levels, 3) increasing the number of *N. crassa*-”preferred” codons does not improve expression of at least two segments of the *cas9* coding region, and 4) *cas9^Hs^*-coding sequences negatively influence transgene expression in a translation dependent manner.

Our data suggest that the relatively large size of a GFP-Cas9 fusion protein should not prevent its detection by our chosen cytological and flow-cytometry-based methods because both methods allowed us to detect a GFP-SAD-6 fusion protein. Importantly, the *gfp-cas9^Hs^* and *gfp-sad-6* transgenes use the same promoter *(ccg-1[P])* and the same *gfp* coding sequences. This suggests that transcription rates of both transgenes should be relatively similar, as should the efficiency of ribosome loading onto the mRNAs derived from each transgene (e.g., because the mRNAs should have identical 5’ UTRs). However, the dissimilarities between the *gfp-cas9^Hs^* and *gfp-sad-6* transgenes are numerous, and they include differences in coding sequences, 3’ UTRs, termination sequences, and locations in the genome. At this point, it is unclear which, if any, of these dissimilar factors are the primary cause of the poor expression of *gfp-cas9^Hs^* in *N. crassa*.

To gain insight into which regions of the *cas9^Hs^* coding sequence may negatively influence its expression, we systematically removed 5’ segments of increasing length from the *cas9^Hs^* coding sequence with a series of *gfp-cas9^HsC#^* transgenes. With this approach, we found it was necessary to remove nearly all the *cas9^Hs^* coding sequence from a *gfp-cas9^HsC#^* transgene before it would express GFP at detectable levels. One possibility is that mRNAs from transgenes with shorter *cas9^Hs^* coding sequences are found at higher levels in *N. crassa* than are mRNAs with longer *cas9^Hs^* coding sequences. If true, this could be due to differences in transcriptional efficiency or transcript stability. However, while we have not measured *gfp-cas9^Hs^* mRNA levels in this study, we did measure GFP levels in strains carrying *gfp-stop-cas9^Hs^* transgenes and we found that all *gfp-stop-cas9^Hs^* transgenes express GFP at detectable levels, even if the transgene contained the full length *cas9^Hs^* coding sequence. These findings suggest that *gfp-cas9^Hs^* expression problems are encountered after *N. crassa* ribosomes finish translating *gfp* sequence and begin translating *cas9^Hs^* sequences.

Codon optimization has long been considered a useful technique for improving the expression of heterologous sequences in transgenic organisms. In this study, we sought to improve the expression of *gfp-cas9^HsC213^* and *gfp-cas9^HsC60^* transgenes by increasing the codon adaptation index (CAI) (Sharp and Li 1987) of each transgene. This codon optimization strategy involves calculating relative adaptiveness (RA) values for each of the 64 possible codons from a set of species-specific high abundance mRNAs and exchanging codons with low RA values for synonymous codons with high RA values. We focused our efforts on *gfp-cas9^HsC213^* and *gfp-cas9^HsC60^* because the *cas9^Hs^* coding sequences in both transgenes are relatively short, and even though *gfp-cas9^HsC60^* expresses GFP, it does so at a level that is an order of magnitude below that of the shortest transgene examined in this study (i.e., *gfp-cas9^HsC4^*). Therefore, there appears to be room to improve the expression of both *gfp-cas9^HsC213^* and *gfp-cas9^HsC60^* transgenes. Interestingly, the *N. crassa* optimized transgenes *gfp-cas9^NcC213^* and *gfp-cas9^NcC60^* did not express GFP at higher levels than their human optimized counterparts (Figure 4). This finding suggests that *N. crassa* codon preferences are neither the major factor preventing expression of *gfp-cas9^HsC213^* at detectable levels, nor the major factor limiting expression of *gfp-cas9^HsC60^*.

A possible explanation of the results observed in this study is that the *N. crassa* translational machinery stalls within the *cas9^Hs^* segments of *cas9^Hs^*-containing mRNAs. To shed light on this possibility, we placed stop codons between *gfp* and *cas9^Hs^* sequences, effectively placing *cas9^Hs^* “coding” sequences in the 3’ UTR of each transgene-derived mRNA. The presence of *cas9^Hs^* coding sequences within the 3’ UTRs of *gfp-stop-cas9^Hs^* transgene derived mRNAs influenced expression in unexpected ways. For example, while transgene expression was detected for all *gfp-siop-cas9^HC#^* transgenes, it was highest for transgenes containing the smallest number of *cas9^Hs^* codons in the 3’ UTR. This finding is consistent with the faux 3’ UTR model (Amrani *et al*. 2006; Nicholson *et al*. 2010; Zhang and Sachs 2015), where abnormally long 3’ UTRs are thought to trigger degradation of the mRNA by nonsense mediated decay. However, our observations are not completely consistent with the faux 3’ UTR model because transgene levels were not strictly directly proportional to the length of the 3’ UTR. For example, transgene expression levels were higher when the 3’ UTR contained between 796 and 612 *cas9^Hs^* codons than when the 3’ UTR contained between 535 and 213 codons. Currently, we do not have an explanation for this phenomenon. More importantly, we performed this set of experiments to determine if halting translation of *gfp-cas9^Hs^* transgenes before ribosomes entered the *cas9^Hs^* regions of an mRNA would improve their expression, and indeed, we found that it does. GFP production was higher from every *gfp-stop-cas9^HsC#^* transgene than from the equivalent *gfp-cas9^Hs^* counterpart, suggesting that the primary obstacle to *cas9^Hs^* expression in *N. crassa* occurs during translation of *cas9^Hs^* sequences.

Our investigation of *cas9^Hs^* expression in *N. crassa* evolved from a desire to study *cas9*-based synthetic gene drivers in filamentous fungi, an area of research that complements our primary focus on fungal meiotic drive elements (Hammond *et al*. 2012; Harvey *et al*. 2014; Pyle *et al*. 2016; Svedberg *et al*. 2018; Rhoades *et al*. 2019a). Although our current findings suggest that *cas9^Hs^* sequences are poorly expressed in *N. crassa*, other researchers have successfully used *cas9^Hs^* to edit genes in *N. crassa* (Matsu-Ura *et al*. 2015). Additionally, Cas9-based technologies have been used with success in dozens of filamentous fungi (Song *et al*. 2019). Still, identifying the mechanistic basis of *cas9^Hs^*’s poor expression in *N. crassa* could help increase the usefulness of Cas9-based technologies in this organism.

## ACKNOWLEDGEMENTS

We would like to thank past and present members of the Hammond Lab for technical assistance with this work. Specifically, we would like to thank ISU undergraduate student Turner Reed for early work on quantifying *gfp-cas9^Hs^* expression levels, as well as ISU undergraduate students Olu Bamidele, Logan Gaskill, Brock Lynn, Jackson Edwards, Trevor Hitzler, and Xhejs Lame for isolating some of the homokaryotic *gfp-cas9^HsC^* strains analyzed in this study. The ISU BDS FACSMelody system was funded by NSF MRI grant 1725199. The ISU Confocal Microscopy Facility was funded by NSF grant DBI-1828136. This work was supported by awards to TMH from the National Science Foundation (1615626 / 2005295).

---

**Figure S1.**
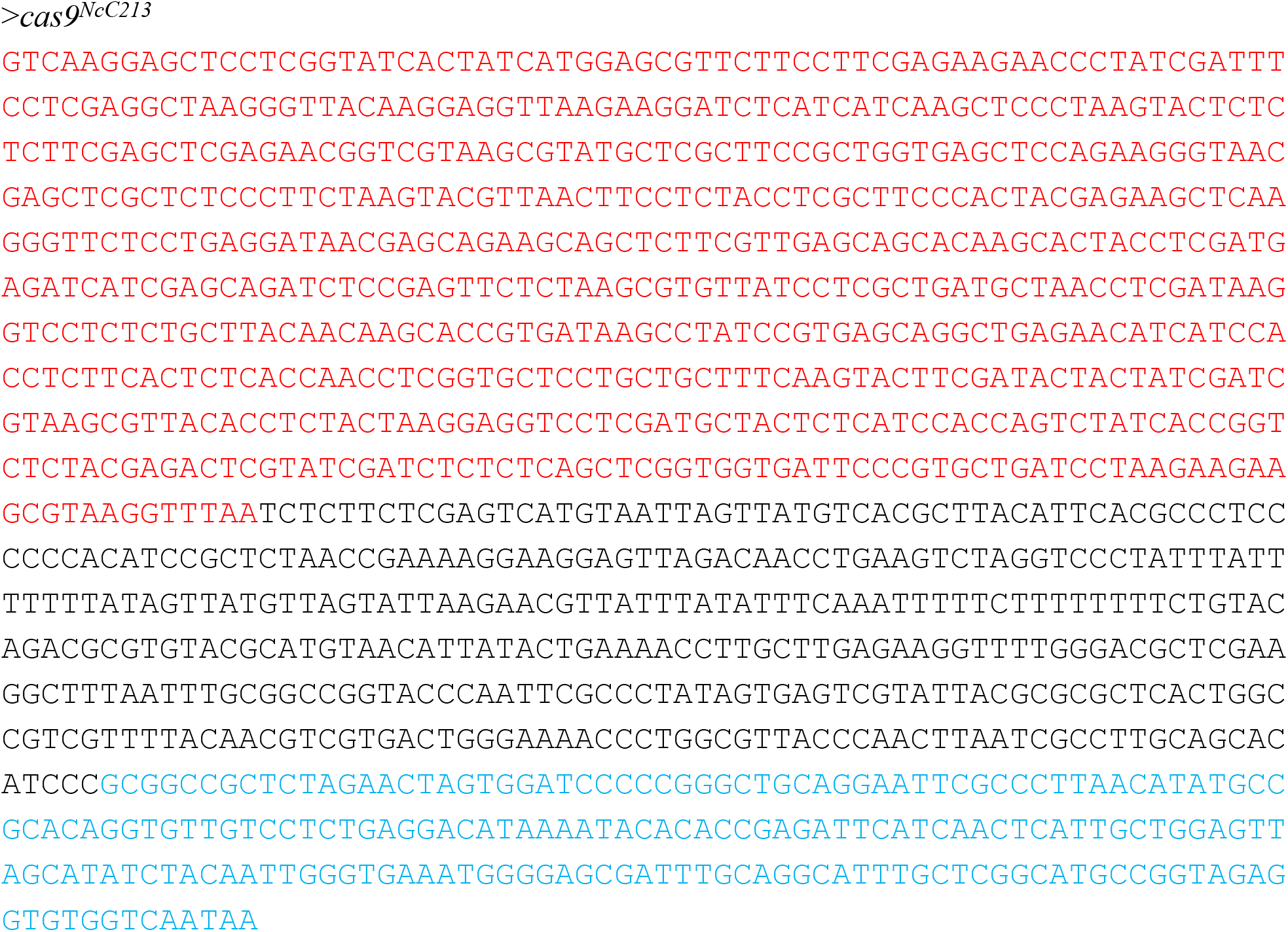
*cas9^NcC213^* sequence. The sequence of *cas9^NcC213^* was ordered as a gBlock^®^ (Integrated DNA technologies) and inserted into pJET1.2 to create plasmid pNR177.2. The *cas9* coding sequence is depicted in red font. There are 221 codons: 213 codons for the *cas9* C terminal end, seven codons for the SV40 NLS, and a single stop codon. The remainder of the sequence includes the *S. cerevisiae cyc-1* terminator (black font) and a pTH1150.1 plasmid sequence to facilitate transformation vector construction (blue font).

**Figure S2.**
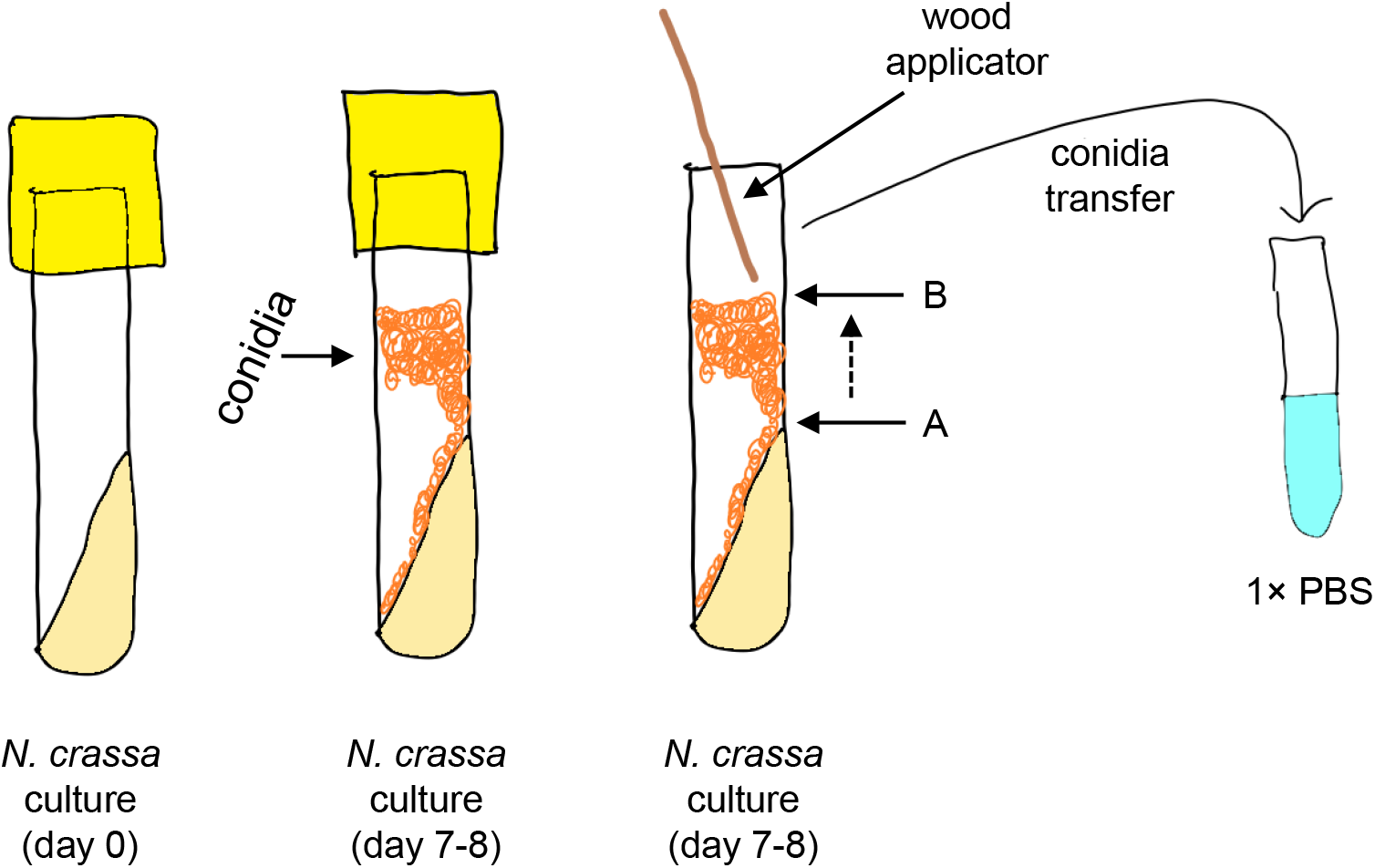
Conidia sampling method. VMM slants were inoculated with *N. crassa* by qualitative transfer of conidia. After a 7–8 day incubation period (see methods), an approximately 2.5 mm path of conidia, from points “A” to “B” in the diagram, were transferred to tubes containing 1× PBS.

**Figure S3.**
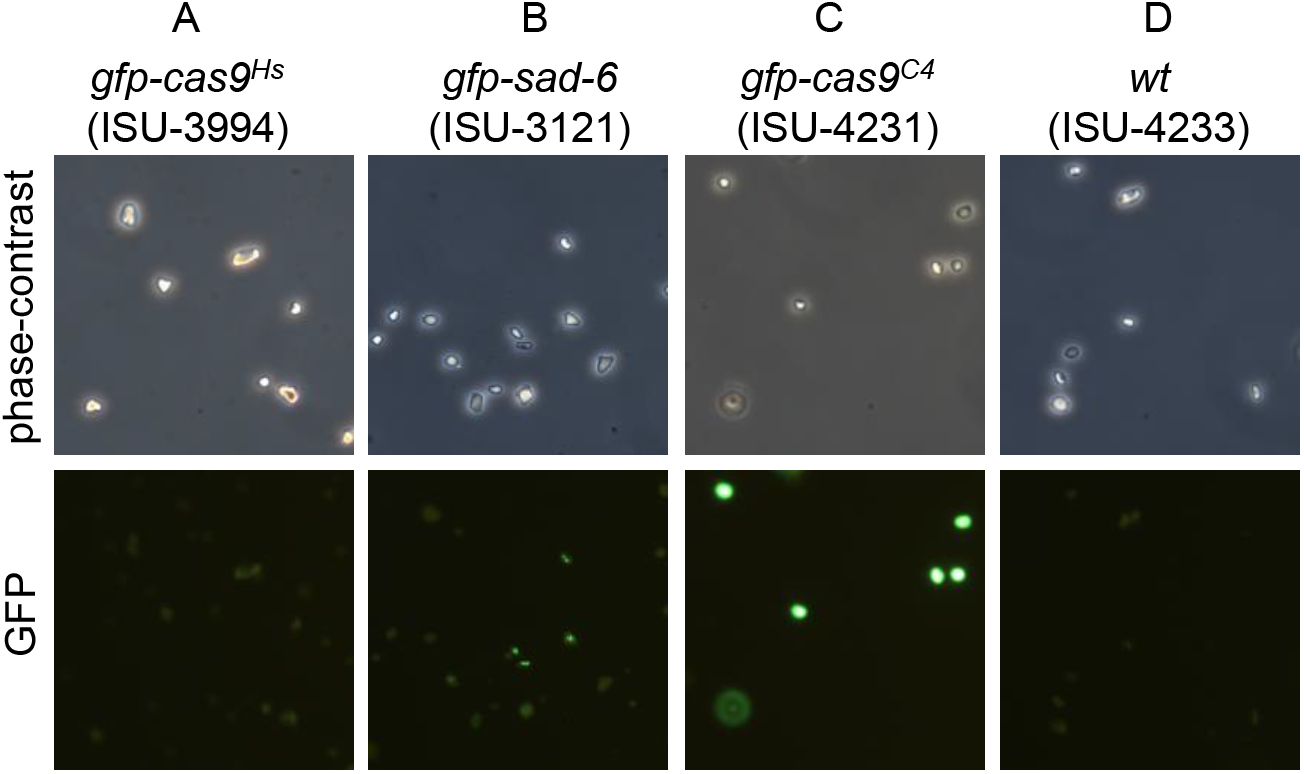
Fluorescence microcopy-based analysis of transgene expression. (A) Conidia from a control strain (ISU-4233), a *gfp-cas9^Hs^* strain (ISU-3994), a *gfp-cas9^HsC4^* strain (ISU-4231), and a *gfp-sad-6* strain (ISU-3121) were examined by fluorescence microscopy with a Leica DMBRE microscope and imaging system.

**Figure S4.**
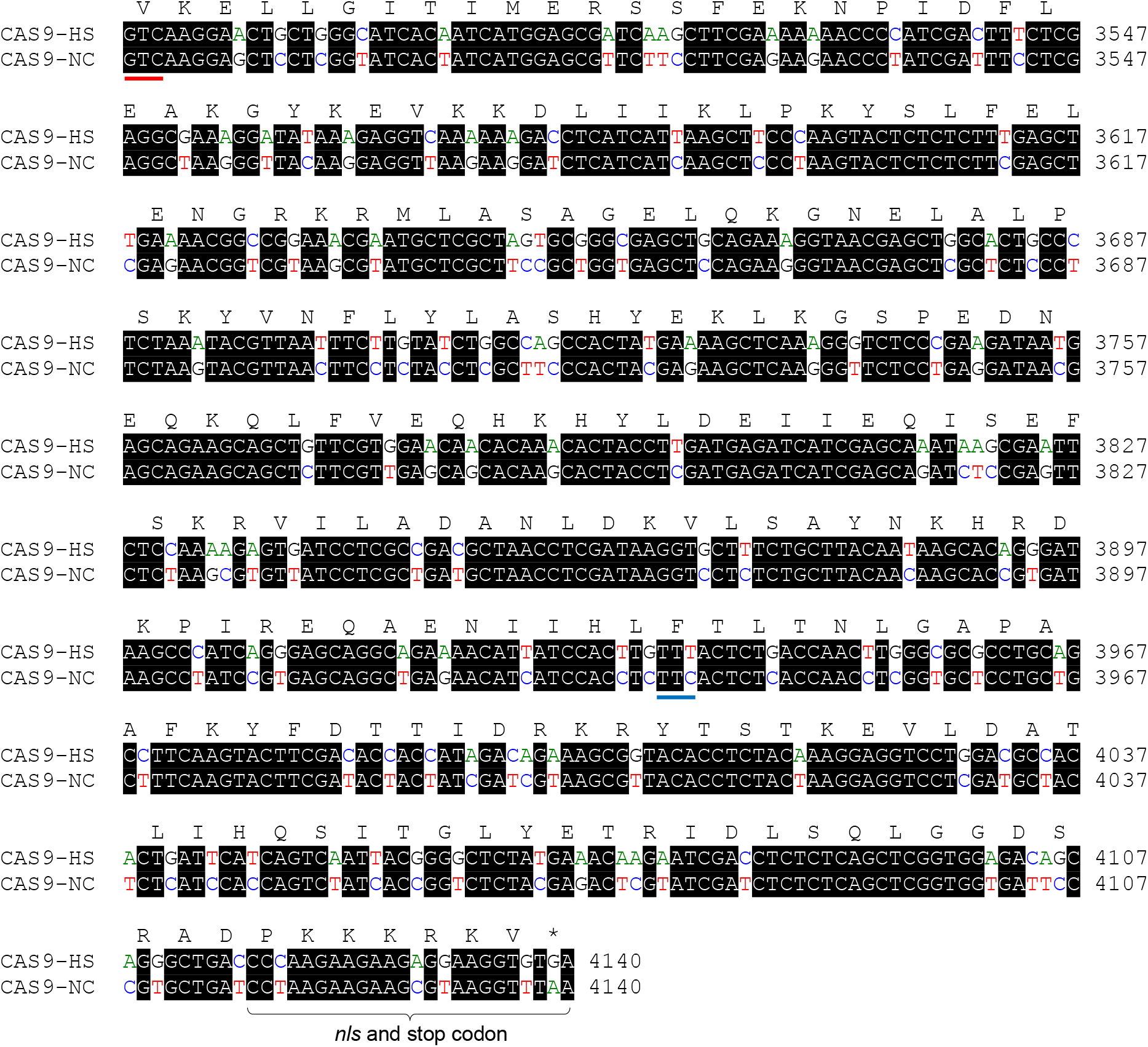
Comparison of *cas9^HsC213^* and *cas9^NcC213^* sequences. A sequence alignment of *cas9^HsC213^* and *cas9^NcC213^* is shown. A total of 120 codons in *cas9^HsC213^* were changed to synonymous codons that are found at higher frequencies in highly expressed *N. crassa* genes. The first codons in the *cas9^Hs60^* and *cas9^Nc60^* are marked with a blue horizontal bar.

**Table S1.**
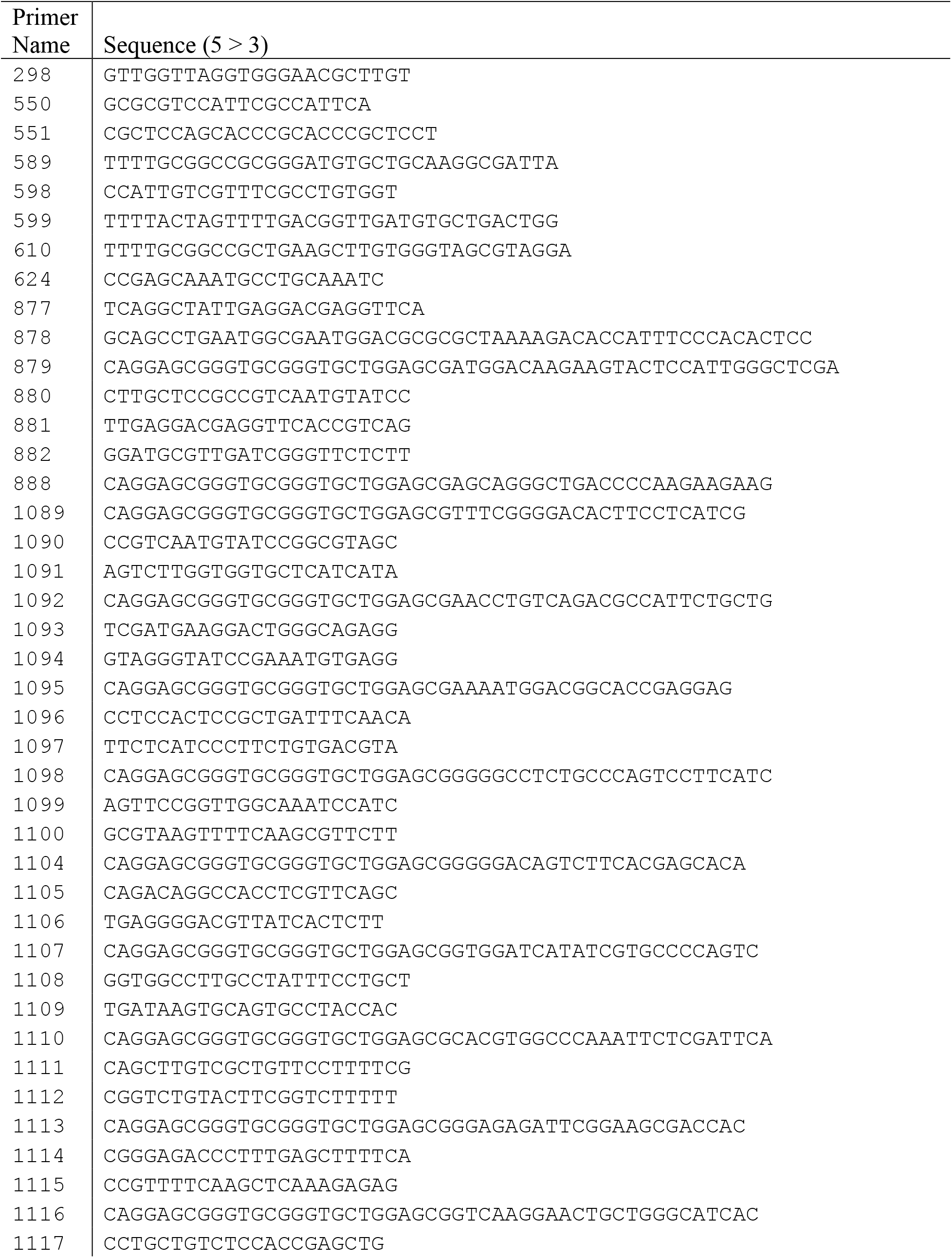
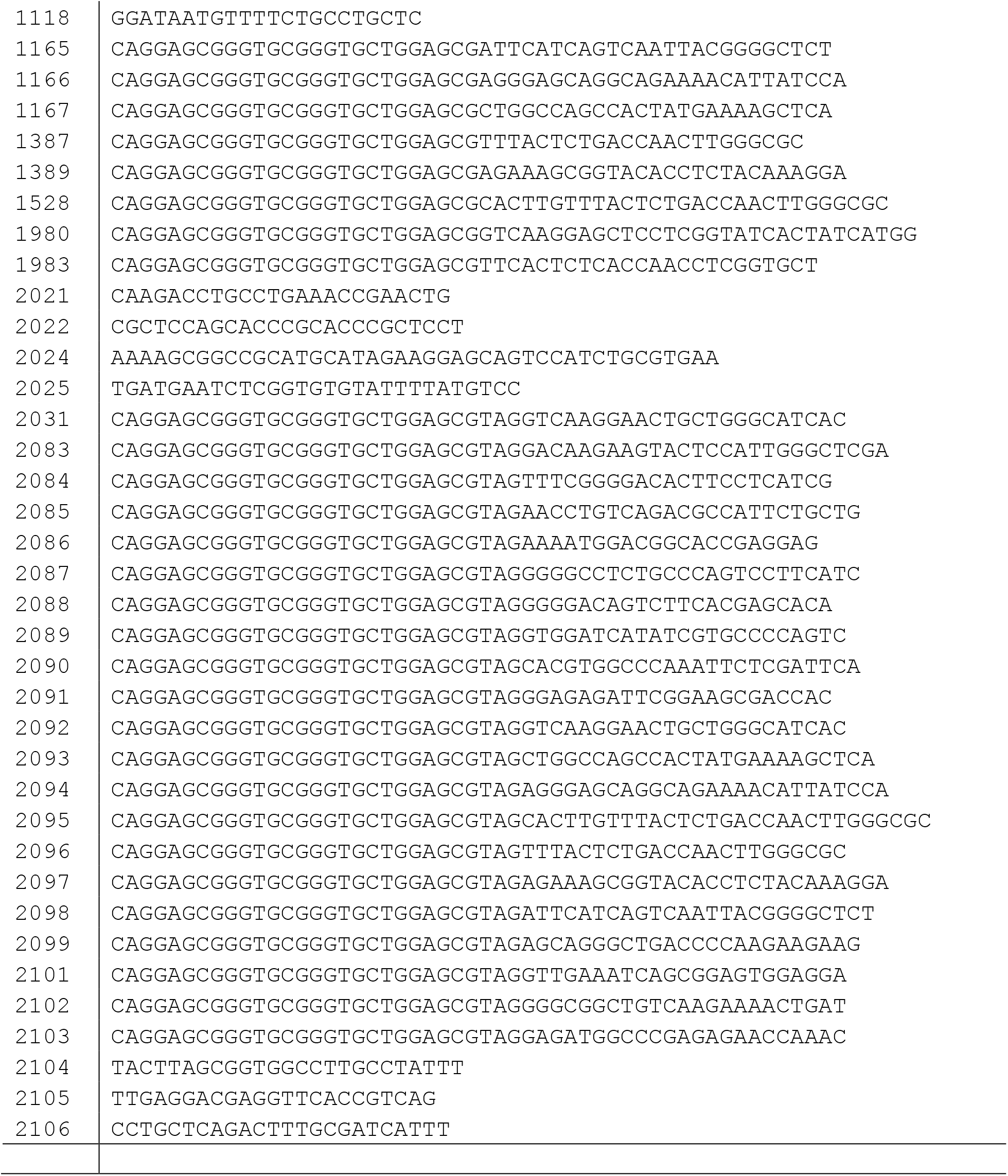
Primers used in this study

**Table S2.** Primer combinations for transformation vector construction by DJ-PCR

| Vector name | Left Fr | Left Rv | Cen Fr | Cen Rv | Right Fr | Right Rv | Nest Fr | Nest Rv |
| --- | --- | --- | --- | --- | --- | --- | --- | --- |
| v76 | 877 | 878 | 550 | 551 | 879 | 880 | 881 | 882 |
| v76-stop | 877 | 878 | 550 | 551 | 2083 | 880 | 881 | 882 |
| v77 | 877 | 878 | 550 | 551 | 888 | 298 | 881 | 624 |
| v77-stop | 877 | 878 | 550 | 551 | 2099 | 298 | 881 | 624 |
| v95 | 877 | 878 | 550 | 551 | 1089 | 1090 | 881 | 1091 |
| v95-stop | 877 | 878 | 550 | 551 | 2084 | 1090 | 881 | 1091 |
| v96 | 877 | 878 | 550 | 551 | 1092 | 1093 | 881 | 1094 |
| v96-stop | 877 | 878 | 550 | 551 | 2085 | 1093 | 881 | 1094 |
| v97 | 877 | 878 | 550 | 551 | 1095 | 1096 | 881 | 1097 |
| v97-stop | 877 | 878 | 550 | 551 | 2086 | 1096 | 881 | 1097 |
| v98 | 877 | 878 | 550 | 551 | 1098 | 1099 | 881 | 1100 |
| v98-stop | 877 | 878 | 550 | 551 | 2087 | 1099 | 881 | 1100 |
| v100 | 877 | 878 | 550 | 551 | 1104 | 1105 | 881 | 1106 |
| v100-stop | 877 | 878 | 550 | 551 | 2088 | 1105 | 881 | 1106 |
| v101 | 877 | 878 | 550 | 551 | 1107 | 1108 | 881 | 1109 |
| v101-stop | 877 | 878 | 550 | 551 | 2089 | 1108 | 881 | 1109 |
| v102 | 877 | 878 | 550 | 551 | 1110 | 1111 | 881 | 1112 |
| v102-stop | 877 | 878 | 550 | 551 | 2090 | 1111 | 881 | 1112 |
| v103 | 877 | 878 | 550 | 551 | 1113 | 1114 | 881 | 1115 |
| v103-stop | 877 | 878 | 550 | 551 | 2091 | 1114 | 881 | 1115 |
| v104 | 877 | 878 | 550 | 551 | 1116 | 1117 | 881 | 1118 |
| v104-stop | 877 | 878 | 550 | 551 | 2092 | 1117 | 881 | 1118 |
| v124 | 877 | 878 | 550 | 551 | 1167 | 298 | 881 | 624 |
| v124-stop | 877 | 878 | 550 | 551 | 2093 | 298 | 881 | 624 |
| v125 | 877 | 878 | 550 | 551 | 1166 | 298 | 881 | 624 |
| v125-stop | 877 | 878 | 550 | 551 | 2094 | 298 | 881 | 624 |
| v126 | 877 | 878 | 550 | 551 | 1165 | 298 | 881 | 624 |
| v126-stop | 877 | 878 | 550 | 551 | 2098 | 298 | 881 | 624 |
| v161 | 877 | 878 | 550 | 551 | 1387 | 298 | 881 | 624 |
| v161-stop | 877 | 878 | 550 | 551 | 2096 | 298 | 881 | 624 |
| v163 | 877 | 878 | 550 | 551 | 1389 | 298 | 881 | 624 |
| v163-stop | 877 | 878 | 550 | 551 | 2097 | 298 | 881 | 624 |
| v193 | 877 | 878 | 550 | 551 | 1528 | 298 | 881 | 624 |
| v193-stop | 877 | 878 | 550 | 551 | 2095 | 298 | 881 | 624 |
| v296-stop | 877 | 878 | 550 | 551 | 2031 | 1117 | 881 | 1118 |
| v307-stop | 877 | 878 | 550 | 551 | 2101 | 2104 | 2105 | 2106 |
| v308-stop | 877 | 878 | 550 | 551 | 2102 | 2104 | 2105 | 2106 |
| v309-stop | 877 | 878 | 550 | 551 | 2103 | 2104 | 2105 | 2106 |

Construction of transformation vectors by DJ-PCR was performed by amplification of left flanks, center fragments, and right flanks with the primers indicated above. For left and right flanks, genomic DNA from strain ISU-4145 was used as the template. For center fragments, plasmid pTH1117.12 (GenBank JF749202.1) was used as the template. After the flanks and center fragments were fused by PCR, the fusion products were amplified with nested primers. Please see Hammond *et al.* (2011) a more complete description of DJ-PCR. Abbreviations are as follows: Left flank forward primer, Left Fr; Left flank reverse primer, Left Rv; Center fragment forward primer, Cen Fr; Center fragment reverse primer, Cen Rv; Right flank forward primer, Right Fr; Right flank reverse primer, Right Rv; Nested forward primer, Nest Fr; and, Nested reverse primer, Nest Rv.

**Table S3.** A set of 100 *N. crassa* high abundance mRNAs

| Rank | Gene | Rank | Gene | Rank | Gene |
| --- | --- | --- | --- | --- | --- |
| 1 | NCU06110T0 | 35 | NCU05804T0 | 69 | NCU00726T0 |
| 2 | NCU01528T0 | 36 | NCU03302T0 | 70 | NCU08964T0 |
| 3 | NCU09345T0 | 37 | NCU03150T0 | 71 | NCU09089T0 |
| 4 | NCU16635T0 | 38 | NCU00618T0 | 72 | NCU07808T0 |
| 5 | NCU00315T0 | 39 | NCU05498T0 | 73 | NCU01546T0 |
| 6 | NCU04553T0 | 40 | NCU04552T0 | 74 | NCU07014T0 |
| 7 | NCU07562T0 | 41 | NCU00979T0 | 75 | NCU01776T0 |
| 8 | NCU05667T0 | 42 | NCU00634T0 | 76 | NCU03148T0 |
| 9 | NCU02003T0 | 43 | NCU02193T0 | 77 | NCU09109T0 |
| 10 | NCU10042T0 | 44 | NCU03102T0 | 78 | NCU07829T0 |
| 11 | NCU05599T0 | 45 | NCU01948T0 | 79 | NCU01949T0 |
| 12 | NCU00294T0 | 46 | NCU06432T0 | 80 | NCU08960T0 |
| 13 | NCU01418T0 | 47 | NCU06431T0 | 81 | NCU07439T0 |
| 14 | NCU03988T0 | 48 | NCU10498T0 | 82 | NCU02181T0 |
| 15 | NCU05561T0 | 49 | NCU09475T0 | 83 | NCU07857T0 |
| 16 | NCU01754T0 | 50 | NCU06047T0 | 84 | NCU00635T0 |
| 17 | NCU00971T0 | 51 | NCU08332T0 | 85 | NCU03757T0 |
| 18 | NCU08389T0 | 52 | NCU00258T0 | 86 | NCU05816T0 |
| 19 | NCU07817T0 | 53 | NCU03753T0 | 87 | NCU03565T0 |
| 20 | NCU09476T0 | 54 | NCU07962T0 | 88 | NCU06226T0 |
| 21 | NCU08963T0 | 55 | NCU07182T0 | 89 | NCU00413T0 |
| 22 | NCU03738T0 | 56 | NCU01962T0 | 90 | NCU08502T0 |
| 23 | NCU08627T0 | 57 | NCU05274T0 | 91 | NCU02437T0 |
| 24 | NCU02250T0 | 58 | NCU04779T0 | 92 | NCU01221T0 |
| 25 | NCU00464T0 | 59 | NCU03703T0 | 93 | NCU16844T0 |
| 26 | NCU01317T0 | 60 | NCU02707T0 | 94 | NCU02744T0 |
| 27 | NCU01827T0 | 61 | NCU01452T0 | 95 | NCU07826T0 |
| 28 | NCU06661T0 | 62 | NCU06892T0 | 96 | NCU05032T0 |
| 29 | NCU03806T0 | 63 | NCU05338T0 | 97 | NCU04114T0 |
| 30 | NCU05554T0 | 64 | NCU09477T0 | 98 | NCU05810T0 |
| 31 | NCU08990T0 | 65 | NCU10069T0 | 99 | NCU06185T0 |
| 32 | NCU00475T0 | 66 | NCU07776T0 | 100 | NCU00706T0 |
| 33 | NCU01552T0 | 67 | NCU06743T0 |  |  |
| 34 | NCU07830T0 | 68 | NCU02905T0 |  |  |

The genes were ranked from highest mRNA abundance (rank 1) to lowest abundance (rank 100).

**Table S4.** Codon RA value estimates for *N. crassa*

| Amino acid | Codon | Total<sup>a</sup> | RA | Amino acid | Codon | Total<sup>a</sup> | RA |
| --- | --- | --- | --- | --- | --- | --- | --- |
| * | TAA | 86 | 1.00 | M | ATG | 366 | 1.00 |
| * | TAG | 9 | 0.10 | N | AAC | 625 | 1.00 |
| * | TGA | 5 | 0.06 | N | AAT | 27 | 0.04 |
| A | GCC | 1044 | 1.00 | P | CCC | 576 | 1.00 |
| A | GCT | 582 | 0.56 | P | CCT | 168 | 0.29 |
| A | GCG | 79 | 0.08 | P | CCG | 46 | 0.08 |
| A | GCA | 22 | 0.02 | P | CCA | 24 | 0.04 |
| C | TGC | 205 | 1.00 | Q | CAG | 527 | 1.00 |
| C | TGT | 10 | 0.05 | Q | CAA | 48 | 0.09 |
| D | GAC | 519 | 1.00 | R | CGC | 726 | 1.00 |
| D | GAT | 234 | 0.45 | R | CGT | 374 | 0.52 |
| E | GAG | 945 | 1.00 | R | AGA | 57 | 0.08 |
| E | GAA | 55 | 0.06 | R | AGG | 44 | 0.06 |
| F | TTC | 491 | 1.00 | R | CGG | 38 | 0.05 |
| F | TTT | 42 | 0.09 | R | CGA | 13 | 0.02 |
| G | GGC | 709 | 1.00 | S | TCC | 567 | 1.00 |
| G | GGT | 583 | 0.82 | S | TCT | 233 | 0.41 |
| G | GGA | 41 | 0.06 | S | AGC | 187 | 0.33 |
| G | GGG | 12 | 0.02 | S | TCG | 119 | 0.21 |
| H | CAC | 362 | 1.00 | S | TCA | 22 | 0.04 |
| H | CAT | 31 | 0.09 | S | AGT | 17 | 0.03 |
| I | ATC | 750 | 1.00 | T | ACC | 722 | 1.00 |
| I | ATT | 165 | 0.22 | T | ACT | 223 | 0.31 |
| I | ATA | 9 | 0.01 | T | ACG | 53 | 0.07 |
| K | AAG | 1563 | 1.00 | T | ACA | 22 | 0.03 |
| K | AAA | 34 | 0.02 | V | GTC | 953 | 1.00 |
| L | CTC | 852 | 1.00 | V | GTT | 285 | 0.30 |
| L | CTT | 249 | 0.29 | V | GTG | 96 | 0.10 |
| L | CTG | 125 | 0.15 | V | GTA | 19 | 0.02 |
| L | TTG | 92 | 0.11 | W | TGG | 158 | 1.00 |
| L | CTA | 7 | 0.01 | Y | TAC | 414 | 1.00 |
| L | TTA | 3 | 0.00 | Y | TAT | 43 | 0.10 |

<sup>a</sup>Total is the total number of occurrences of each codon in the coding sequences of the 100 genes listed in Table S3. Relative adaptiveness (RA) values were calculated according to the method of Sharp and Li (1987).

**Table S5.** MFI values

| Data set<sup>a</sup> | Strain Name | Vector | *cas9* codons remaining | *gfp* fusion position on *cas9* | MFI 7-day culture | MFI 8-day culture |
| --- | --- | --- | --- | --- | --- | --- |
| all | ISU-3121 | na | na (PC) | na (PC) | 435.66 | 454.47 |
| all | ISU-3886 | na | na (NC) | na (NC) | 104.91 | 97.38 |
| all | ISU-4145 | na | na (NC) | na (NC) | 133.15 | 93.09 |
| all | ISU-4233 | an | na (NC) | na (NC) | 112.09 | 102.5 |
| cas | ISU-3994 | v76 | 1372 | 1 | 122.36 | 92.07 |
| cas | ISU-4888 | v95 | 1209 | 164 | 107.42 | 108.21 |
| cas | ISU-4889 | v96 | 1078 | 295 | 107.62 | 97.36 |
| cas | ISU-4092 | v97 | 991 | 382 | 113.54 | 93.36 |
| cas | ISU-4892 | v98 | 888 | 485 | 123.31 | 104.17 |
| cas | ISU-4894 | v100 | 656 | 717 | 126.82 | 99.05 |
| cas | ISU-4895 | v101 | 535 | 838 | 130.97 | 102.76 |
| cas | ISU-4897 | v102 | 444 | 930 | 117.02 | 86.9 |
| cas | ISU-4107 | v103 | 318 | 1055 | 115.71 | 93.82 |
| cas | ISU-4232 | v104 | 213 | 1160 | 125.3 | 108.62 |
| cas | ISU-4234 | v124 | 135 | 1238 | 171.28 | 140.2 |
| cas | ISU-4235 | v125 | 70 | 1303 | 220.01 | 234.57 |
| cas | ISU-4522 | v193 | 62 | 1311 | 511.21 | 370.54 |
| cas | ISU-4618 | v161 | 60 | 1313 | 2251.61 | 1121.58 |
| cas | ISU-4624 | v163 | 40 | 1333 | 5228.12 | 4119.94 |
| cas | ISU-4236 | v126 | 25 | 1349 | 16022.36 | 11923.71 |
| cas | ISU-4231 | v77 | 4 | 1369 | 23894.7 | 19467.86 |
| opt | ISU-4884 | v293 | 213 | 1160 | 94.59 | 100.71 |
| opt | ISU-4886 | v294 | 60 | 1313 | 601.74 | 436.47 |
| stop | ISU-4933 | v76-stop | 1372 | 1 | 260.68 | 198.29 |
| stop | ISU-4942 | v95-stop | 1209 | 164 | 300.98 | 197.57 |
| stop | ISU-4943 | v96-stop | 1078 | 295 | 197.04 | 171.94 |
| stop | ISU-4944 | v97-stop | 991 | 382 | 413.81 | 441.81 |
| stop | ISU-4945 | v98-stop | 888 | 485 | 318.96 | 270.13 |
| stop | ISU-4951 | V307-stop | 796 | 577 | 813.71 | 623.85 |
| stop | ISU-4952 | V308-stop | 714 | 659 | 1262.2 | 688.16 |
| stop | ISU-4946 | v100-stop | 656 | 717 | 945.56 | 906.43 |
| stop | ISU-4953 | v309-stop | 612 | 761 | 2698.72 | 1175.61 |
| stop | ISU-4947 | v101-stop | 535 | 838 | 381.04 | 443.96 |
| stop | ISU-4948 | v102-stop | 443 | 930 | 325.51 | 381.75 |
| stop | ISU-4949 | v103-stop | 318 | 1055 | 347.14 | 215.91 |
| stop | ISU-4875 | v296-stop | 213 | 1160 | 486.69 | 473.05 |
| stop | ISU-4935 | v104-stop | 213 | 1160 | 482.28 | 447.27 |
| stop | ISU-4936 | v124-stop | 135 | 1238 | 4930.99 | 3809.22 |
| stop | ISU-4938 | v193-stop | 62 | 1311 | 3208.58 | 2014.42 |
| stop | ISU-4939 | v161-stop | 60 | 1313 | 19900.07 | 13804.47 |
| stop | ISU-4941 | v163-stop | 40 | 1333 | 20585.8 | 12921.51 |
| stop | ISU-4937 | v126-stop | 25 | 1348 | 25380.18 | 17090.45 |
| stop | ISU-4934 | v77-stop | 4 | 1369 | 44774 | 34718.37 |

<sup>a</sup>These data were used to construct the charts presented in Figure 3 and Figure 4. Dataset: all, strains provided the positive and negative control values in all charts; cas9, strains provided the values used to produce the chart in Figure 3A and Figure 4; stop, strains provided the values used to produce the chart in Figure 3B. Abbreviations: na, not applicable, PC, positive control, GFP-SAD-6 strain; NC, negative control, no GFP transgenes present.

